# The effect of *Stachys inflata* on the performance of broiler chickens and Newcastle Disease Antibody Titer

**DOI:** 10.1101/2021.06.25.449908

**Authors:** Hamidreza Attaran, Abdolkarim Zamanimoghaddam, Alireza Ghannadi, Abdolrasool Namjoo, Farshad Zamani

**Author notes:** Corresponding author: Dr. Hamidreza Attaran, Cell and Molecular Biology and Microbiology Department, Biological Science and Technology College, University of Isfahan, Isfahan, Iran.

## Abstract

This study shows how performance and NDHI titers of broiler chickens are affected by the plant *Stachys inflata*y. One hundred and eighty, one day old Ross male broiler chickens were randomly divided into 5 groups of 36 (consisting of 3 replicates) and placed in 12 separate pens. Groups 1, 2, 3, 4 and 5 received 0, 0.15, 0.3, 0.45 and 0.6% *S*.*inflata* in their feed respectively from 8 to 42 days of age. Performance indexes were recorded on a weekly basis. At 42 days of age, serum and mid gut samples were taken for NDHI test and histopathological studies. The results were analyzed by the One Way ANOVA, Tukey test. The body weight of the second group was significantly (P ≤ 0.05) higher than the control from the third week to the end. All treatment groups had HI titers higher than the control but the differences were not significant (P > 0.05). In the third group (0.3% *S*.*inflata*), relative liver and heart weight were significantly lower than control. No microscopic lesions were observed in mid gut in all experimental groups. It seems that *S. inflata* reduces harmful intestinal flora which causes better use of feed.

## Introduction

The use of natural products is growing in the world, especially in developing countries such as China, India and Arabic countries. Parallel to the use of medicinal herbs in the treatment of human diseases, plants or plant originated compound have been used extensively within the veterinary field especially as natural growth promoters or feed supplement in the broiler industry.

One of the most important medicinal herb families is the *Lamiaceae*, whose related drugs have been used at times for different purposes. According to Skaltsa et al. the sub cosmopolitan genus known as *Stachys*, which is compromised of more than 270 species and is justifiably considered one of the largest genera of the *Lamiaceae* (Skaltsa et al., 2001). For several centuries, some *Stachys* species have been used as herbs for their potential health benefits (Zargari, 1990) and this would include the 34 species which are found in Iran (Mozaffarian, 1996). Along with many other chemicals, Phytochemical lab tests on *Stachys* species have shown presence of phenylethanoid glycosides (Nishimura *et al*., 1991; Miyase et al., 1996), terpenoids, steroids (Ross and Zinchenko, 1975; Yamamoto et al., 1994), diterpenes (Piozzi et al., 1980) and flavonoids (Zinchenko, 1970; EL-Ansari et al., 1991) along with some essential oils of some species (Skaltsa et al., 1999; CËakir et al., 1997). Pharmacological studies have shown that extracts of some *Stachys* species have anti-inflammatory, anti-toxic (Maleki et al., 2001; Zinchenko et al., 1981), anti-nephritic (>Hayashi et al., 1994a, 1994b), anti-hepatitis (Savchenko and Khvorostinka, 1978) and anti-anoxia (Yamahara et al., 1990) properties. Phenylethanoid glycosides, triterpenoids and flavonoids are considered the active components responsible for biological reactions in the *Stachys* genus (Miyase et al., 1996; Yamamoto et al., 1994; Zinchenko, 1970; EL-Ansari et al., 1991).

Related to this genus, the *S. inflata* is one of the known plants in which extracts have been used in Iranian folk medicine as treatment for infective, rheumatic and other inflammatory disorders (Maleki et al., 2001). Several pharmaceutical effects have been proven for *S. inflata* related compounds and its different fractions in vitro and in vivo. For this reason, a study was designed to investigate the beneficial effects of this herb on the immune system by checking Newcastle disease (ND) antibody titer and performance of broiler chickens.

## Material and methods

In this study, 180 day old male Ross 308 broiler chickens were randomly divided into 5 groups of 36 with three replicates of 12. The feed rations were formulated according to the standard of NRC. The flowering aerial parts of *S. inflata* including stems and leaves have been collected during the flowering period (June 2009), identified in Central Herbarium of Iran (Research Institute of Forest and Rangelands, Tehran). *S. inflata* were collected from Khalkhal, in the province of Ardabil, Iran. Dried plant materials were grounded to fine powder. Groups 2, 3, 4 and 5 received a balanced ration with 0.15%, 0.3%, 0.45% and 0.6% of *S. inflata* respectively. Group 1 as the control, received a balanced diet without any *S. inflata* added to their feed until the end of the 42 days experiment. Chicks received dietary treatments at free access from 1 to 42 days of age. All chicks were vaccinated against ND, Infectious bronchitis (IB) and Infectious bursal disease (IBD) according to the regional standard vaccination program. All other management procedures were the same for all groups. The following parameters were recorded on a weekly basis: feed intake, mean body weight, feed conversion ratio (FCR) and mortality rate. Processing criteria obtained was body weight, liver weight, heart weight, gizzard weight, breast weight, thigh weight and abdominal fat weight. Four chicks from each pen were randomly selected and blood samples were collected for NDHI tests at 31 and 42 days of age (10 days after ND vaccination). Mid-guts of sacrificed birds of all groups were examined for histopathological changes. Other special conditions such as clinical signs and diarrhea index (by visiting the birds 3 times a day to access the probable incidence of diarrhea) were investigated throughout the experiment.

### Statistical analysis

All data were analyzed by One Way ANOVA and in case of a significant difference also by Tukey’s test.

## Results

### Effects of *S. inflata* on Body Weight

At the end of the 3^rd^, 4^th^, 5^th^ and 6^th^ weeks, the mean body weight of group 2 (0.15%) was significantly higher than the control group (P ≤ 0.05). In the 6^th^ week, also among group 2 with groups 4 and 5, a significant difference was observed. At the end of the second week, weight gain had been more in all treatment groups than in the control group, but significant differences between any of the groups were not observed. At the 3^rd^, 4^th^ and 5^th^ weeks the statistical significant difference between the groups receiving 0.15 percent and the control group at the level of P ≤ 0.05 was observed. At the 6^th^ week the significant difference was observed between 0.15 percent recipient group among 0.45 and 0.6 percent recipient groups at the level of P ≤ 0.05.

### Effect of *S. inflata* on feed intake (FI)

No significant difference on feed intake between groups was observed (P > 0.05). The feed consumption during 1-42 days was approximately the same in all groups, and there was no significant difference at the level of P ≤ 0.05 in different groups (Table 2).

**Table 1.**
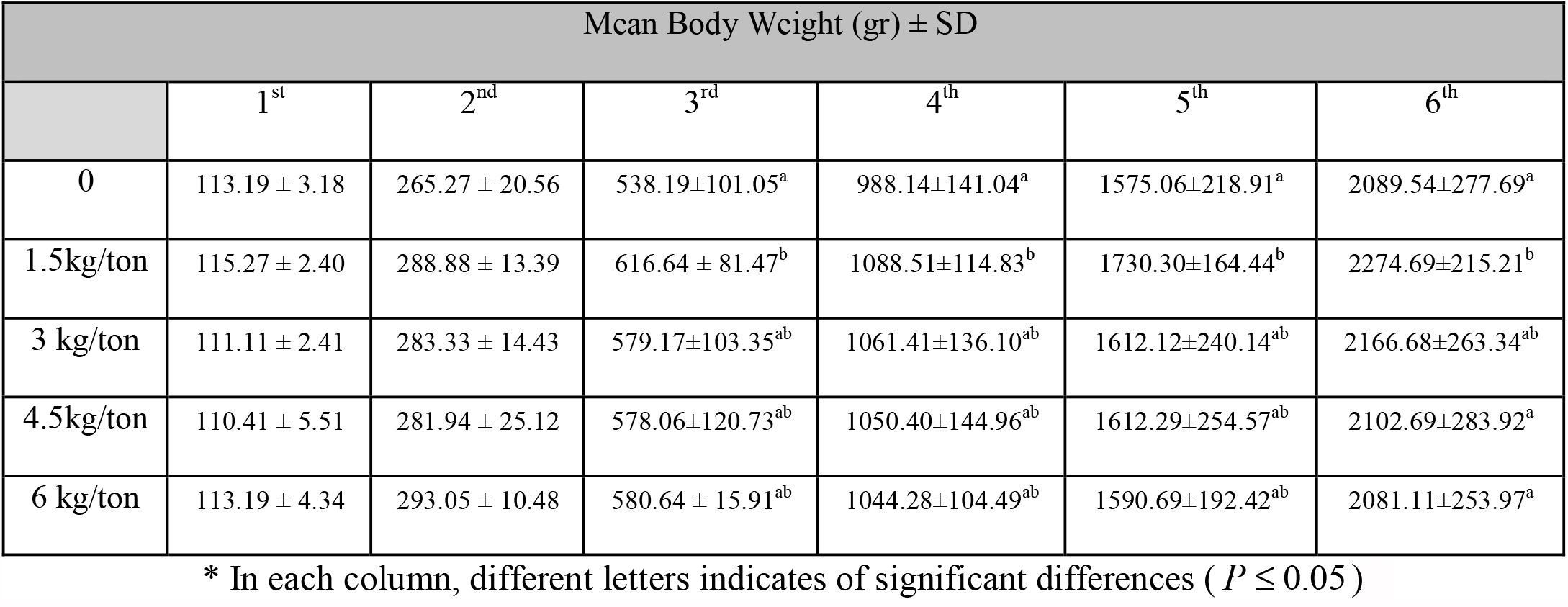
Effect of *S.inflata* on Body Weight.

| Mean Body Weight (gr) ± SD |  |  |  |  |  |  |
| --- | --- | --- | --- | --- | --- | --- |
|  | 1<sup>st</sup> | 2<sup>nd</sup> | 3<sup>rd</sup> | 4<sup>th</sup> | 5<sup>th</sup> | 6<sup>th</sup> |
| 0 | 113.19 ± 3.18 | 265.27 ± 20.56 | 538.19±101.05<sup>a</sup> | 988.14±141.04<sup>a</sup> | 1575.06±218.91<sup>a</sup> | 2089.54±277.69<sup>a</sup> |
| 1.5kg/ton | 115.27 ± 2.40 | 288.88 ± 13.39 | 616.64 ± 81.47<sup>b</sup> | 1088.51±114.83<sup>b</sup> | 1730.30±164.44<sup>b</sup> | 2274.69±215.21<sup>b</sup> |
| 3 kg/ton | 111.11 ± 2.41 | 283.33 ± 14.43 | 579.17±103.35<sup>ab</sup> | 1061.41±136.10<sup>ab</sup> | 1612.12±240.14<sup>ab</sup> | 2166.68±263.34<sup>ab</sup> |
| 4.5kg/ton | 110.41 ± 5.51 | 281.94 ± 25.12 | 578.06±120.73<sup>ab</sup> | 1050.40±144.96<sup>ab</sup> | 1612.29±254.57<sup>ab</sup> | 2102.69±283.92<sup>a</sup> |
| 6 kg/ton | 113.19 ± 4.34 | 293.05 ± 10.48 | 580.64 ± 15.91<sup>ab</sup> | 1044.28±104.49<sup>ab</sup> | 1590.69±192.42<sup>ab</sup> | 2081.11±253.97<sup>a</sup> |

\* In each column, different letters indicates of significant differences ( $P \leq 0.05$ )

**Table 2.**
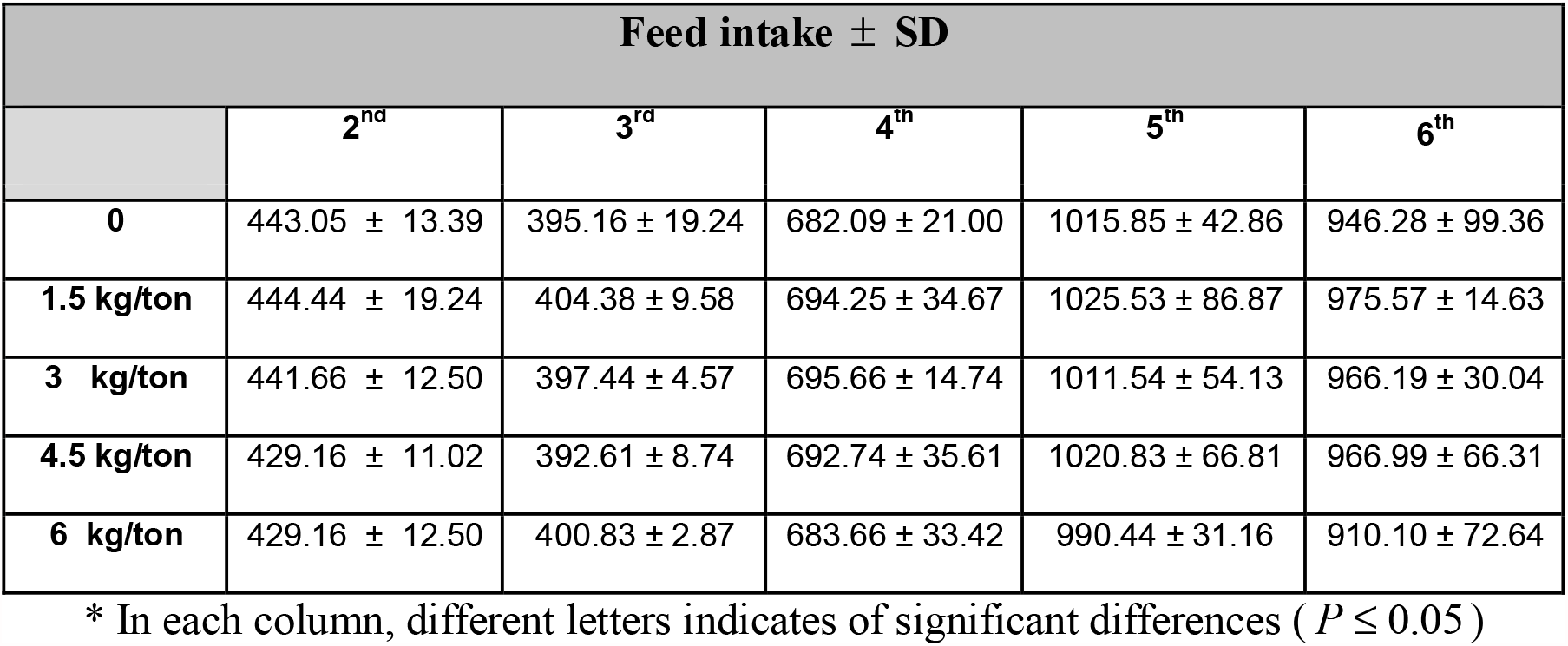
Effect of *S.inflata* on Feed intake (FI)

### Effect of *S. inflata* on feed conversion ratio (FCR)

Feed conversion ratio at 3 weeks of age was significantly improved when 0.15% of *S. inflata* was added to the control diet. According to table 3, observation showed that FCR at 21 days old chicks in the 0.15 percent received group was significantly (P ≤ 0.05) less than in the control group. In the rest of the growing period 21 days through to 42 days, the 0.15 percent recipient group had a lower FCR, but there were no significant differences (P > 0.05).

**Table 3.** Effect of *S.inflata* on Feed Conversion Ratio (FCR)

| Feed Conversion Ratio ± SD | | | | | |
| --- | --- | --- | --- | --- | --- |
| F.C.R | 3<sup>rd</sup> | 4<sup>th</sup> | 5<sup>th</sup> | 6<sup>th</sup> | Total FCR |
| 0 | 1.63 ± 0.1<sup>a</sup> | 1.59 ± 0.03 | 1.68 ± 0.09 | 1.81 ± 0.01 | 1.68 ± 0.01 |
| 1.5 kg/ton | 1.41 ± 0.02<sup>b</sup> | 1.52 ± 0.03 | 1.60 ± 0.03 | 1.73 ± 0.01 | 1.57 ± 0.14 |
| 3 kg/ton | 1.53 ± 0.01<sup>ab</sup> | 1.53 ± 0.07 | 1.66 ± 0.02 | 1.82 ± 0.08 | 1.64 ± 0.14 |
| 4.5 kg/ton | 1.57 ± 0.08<sup>ab</sup> | 1.54 ± 0.07 | 1.69 ± 0.07 | 1.98 ± 0.24 | 1.69 ± 0.20 |
| 6 kg/ton | 1.49 ± 0.05<sup>ab</sup> | 1.61 ± 0.15 | 1.69 ± 0.03 | 1.98 ± 0.25 | 1.69 ± 0.21 |

\* In each column, different letters indicates of significant differences ( $P \leq 0.05$ )

### Effects of *S. inflata* on NDHI titers

Newcastle antibody titers were also measured in two stages, once at day 31 and again at day 42 days where the results are expressed in table 4. According to these results, all groups which used *S. inflata* in their premixes had greater antibody titers than the control group at these two stages. No significant differences were observed in any of these groups (P > 0.05).

**Table 4.**
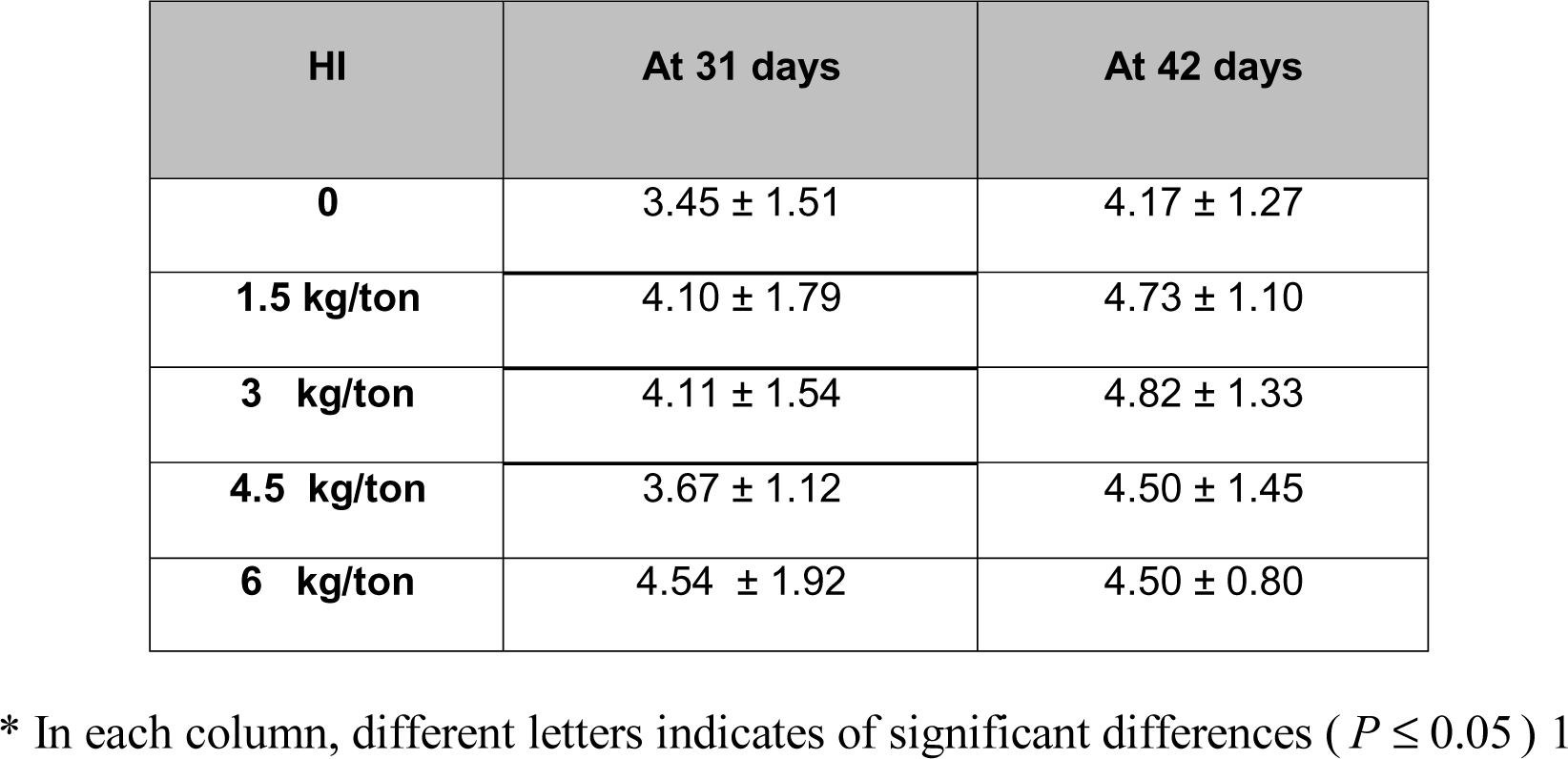
Effect of *S.inflata* on NDHI titer.

| HI | At 31 days | At 42 days |
| --- | --- | --- |
| 0 | 3.45 ± 1.51 | 4.17 ± 1.27 |
| 1.5 kg/ton | 4.10 ± 1.79 | 4.73 ± 1.10 |
| 3 kg/ton | 4.11 ± 1.54 | 4.82 ± 1.33 |
| 4.5 kg/ton | 3.67 ± 1.12 | 4.50 ± 1.45 |
| 6 kg/ton | 4.54 ± 1.92 | 4.50 ± 0.80 |

\* In each column, different letters indicates of significant differences ( $P \leq 0.05$ ) 1

### Effect of *S* .*inflata* on the relative weight of breast, thigh muscles, liver, gizzard, heart, abdominal and gizzard fat

As it is observed in table 5, the percentage of breast’s weight and thigh muscle’s weight between the groups were not significant (P > 0.05). Treatment groups 2 and 3 with 0.15% and 0.3% *S. inflata* had the lowest liver weight respectively. Between groups 3 with 0.3% *S. inflata* and the control group, there is a relatively noticeable liver weight difference (P ≤ 0.05). The gizzard’s weight from the groups that were treated showed no differences (P > 0.05). The highest and lowest heart weight was related to the control and 0.3% treatment group respectively (P ≤ 0.05). No significant difference was observed among the experimental groups in abdominal and gizzard fat (P > 0.05).

**Table 5.** Effect of *S.inflata* on the relative weight of breast and thigh muscles, liver, gizzard, heart, abdominal and gizzard fat.

| the relative weight ± SD | | | | | | | |
| --- | --- | --- | --- | --- | --- | --- | --- |
|  | Breast | Thigh | Liver | Gizzard | Heart | Abdominal fat | Gizzard fat |
| **0** | 21.00 ± 2.13 | 21.00 ± 1.17 | 2.39± 0.23<sup>a</sup> | 1.60 ± 0.23 | 0.46± 0.05<sup>a</sup> | 0.88 ± 0.29 | 0.48 ± 0.13 |
| **1.5 kg/ton** | 21.30 ± 0.73 | 21.68 ± 1.15 | 1.90±0.23<sup>ab</sup> | 1.58 ± 0.13 | 0.43±0.05<sup>ab</sup> | 0.89 ± 0.38 | 0.53 ± 0.34 |
| **3 kg/ton** | 21.51 ± 0.60 | 21.36 ± 0.46 | 1.81± 0.56<sup>b</sup> | 1.31 ± 0.17 | 0.33± 0.11<sup>b</sup> | 0.92 ± 0.55 | 0.29 ± 0.06 |
| **4.5 kg/ton** | 20.67 ± 0.74 | 20.99 ± 0.73 | 2.22±0.26<sup>ab</sup> | 1.52 ± 0.15 | 0.42±0.05<sup>ab</sup> | 0.94 ± 0.30 | 0.60 ± 0.31 |
| **6 kg/ton** | 21.60 ± 0.68 | 20.76 ± 0.80 | 2.14±0.23<sup>ab</sup> | 1.51 ± 0.17 | 0.43±0.06<sup>ab</sup> | 1.06 ± 0.16 | 0.46 ± 0.31 |

\* In each column, different letters indicates of significant differences ( $P \leq 0.05$ ) 4

### Effect of *S. inflata* on pathological examination in mid gut

No microscopic lesions were discovered in the mid-gut of all the groups. Observations show that the lowest diameter of mid gut was related to 0.45% and 0.15% groups respectively but no significant difference was observed between the groups (P > 0.05). Also between the diameter of mucosal and sub-mucosal layer, the lowest measurement of mucosal and sub-mucosal layer of mid-gut belongs to the 0.45% group. (Table 6)

**Table 6.**
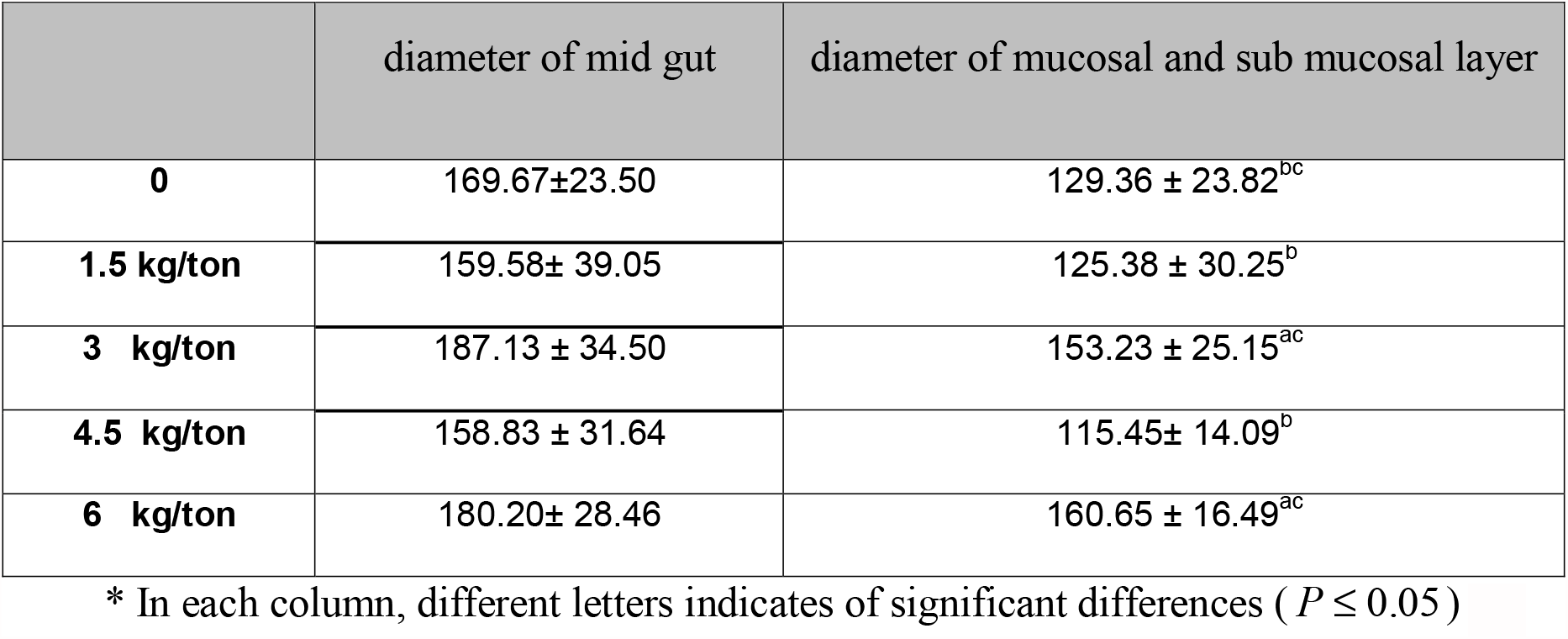
Effect of *Stachys inflata* on pathological examination in mid gut.

## Discussion

Body weight gains by using *S. inflata* could be in relation to higher anti-oxidants, anti-microbial properities, and anti-inflammatory properties. It seems that *S. inflata* reduces harmful intestinal flora which causes better use of feed. On the other hand, *S. inflata* can prevent the feed from harmful effects of mycotoxins. Furthermore, *S. inflata* with its high anti-inflammatory properties, can reduce layers of intestinal inflammation and lead to better feed absorption. Anti-oxidant activities of the plant may also prevent and protect the chickens ration from oxidation and save the value of vitamins and proteins of the diet.

There was no significant difference on feed intake among the treatment groups and the control group (P > 0.05). Lack of significant differences between the control group and all of the groups receiving *S* .*inflata* showed that improvement in overall weight in the recipient groups is not due to higher feed consumption, but due to the better use of food components and increase efficiency of the feed. It could be a postulated improvement in FCR by reducing inflammation and harmful intestinal bacteria and improve the feed absorption from the gastro-intestinal tract.

Although there were no significant differences in NDHI observed in any of the groups (P > 0.05), antibody titers in all groups which had used *S. inflata* as a premix in ration was greater than the control groups antibody titer. Since stress and sickness are two factors that cause vaccination failure, perhaps sedative, anti-inflammatory and anti-microbial properties of this plant could protect chickens against pathogens and stress.

Considering that the relative weight of carcasses, breasts and thigh muscles are mainly affected by genetic factors, inclusion of the feed with *S. inflata* cannot significantly alter these relative weights of thigh muscles and breasts simultaneously.

There was no significant difference between the abdominal fat and the fat around the gizzard of treatment groups and the control group (P > 0.05). This could be (evidence) postulated that adding S. *inflata* to the feed has no detrimental effect on the abdominal fat metabolism.

In 2007, the effect of *Satureja hortensis (S. h.)* on performance of broiler chickens and NDHI titers were investigated. The results showed birds that received 0.3% *S. h*. had a higher body weight and lower FCR, but the differences were not significant (P ≥ 0.05). All groups that received *S. h*. in ration had higher HI titers than the control group. Adding *S. h*. to the diet had no effect on the relative weight of breasts, thigh muscles, abdominal fat and histological structures of the liver and kidney samples (Zamanimoghaddam et al., 2007).

Dulger and Gonuz (2004., 2005) reported the antimicrobial activity of some endemic *Stachys* species which includes *S. sivasica, S. anamurensis, S. cydnia, S. aleurites* and *S. pinardii*. In 2008, research was done on the antimicrobial activities of the methanol extracts from dried flowering aerial parts of *S. byzantina, S. inflata, S. lavandulifolia* and *S. laxa* (Labiatae). According to this research, “Flavonoids may be responsible for their antibacterial activity,” which is extracted from the aerial parts of the genus; Stacchys (Saeedi et al., 2008).

Hydro-alcoholic extract of *S. inflata* has been demonstrated to exert potent anti-inflammatory effects in peripheral inflammation which is induced by carrageenan or formalin in rat paw (Maleki et all., 2001). In another study, the Hydro-alcoholic extracts of *S. inflata* reduced infarct size with a low dosage. This corresponds with the findings of Maleki and co-workers (2001) who revealed that the higher dose of the extract had no anti-inflammatory effects. This result suggests that hydro-alcoholic extracts from aerial parts of *S.inflata* may play a major role in reducing the infarct size (Garjani et al., 2001).

In other studies, there were anti-inflammatory effects of hydro-alcoholic and ethyl acetate extracts from *S. schtschegleevii* in both the flowering and sterile tops (Nazemiyeh et al., 2007; Maleki et al., 2008). The anti-inflammatory and anti-nociceptive properties of total methanolic extracts of the flowering aerial parts of two *Stachys* species (Rezazadeh et al., 2005). Anti-inflammatory and anti-hyperalgesic activities of *Stachys athorecalyx* extracts on CFA-induced Inflammation (Rezazadeh et al., 2009) were also observed.

The stabilizing effects of methanolic extract of *S. byzantina, S. inflata* and *S. laxa* on sunflower oil as antioxidant agents were observed by Morteza-Semnani et al., 2006 and the results indicated that *S. laxa* had a potential source of antioxidants. In 2009, *S. persica* and *S. fruticulosa* revealed anti-oxidant effects approximately two times greater than *S. laxa* (Khanavi et al., 2009). There were also many other studies on the anti-oxidant effects of *S. inflata, S. byzantina, S. setifera* and *S. laxa* (Erdemoglu et al., 2006; Morteza-Semnani et al., 2006). However, because various methods have been used, the results are not fully comparable.

